## Supplementary Information for "Structure of γ-secretase (PSEN1/APH-1B) in complex with Aβ46 provides insights into amyloid-β processing and modulation by the APH-1B isoform"

Ivica Odorcic[1,2,3,4], Mohamed Belal Hamed Soliman[3,4], Sam Lismont[3,4], Lucia Chavez Gutierrez[3,4*$], and Rouslan G. Efremov[1,2*$]

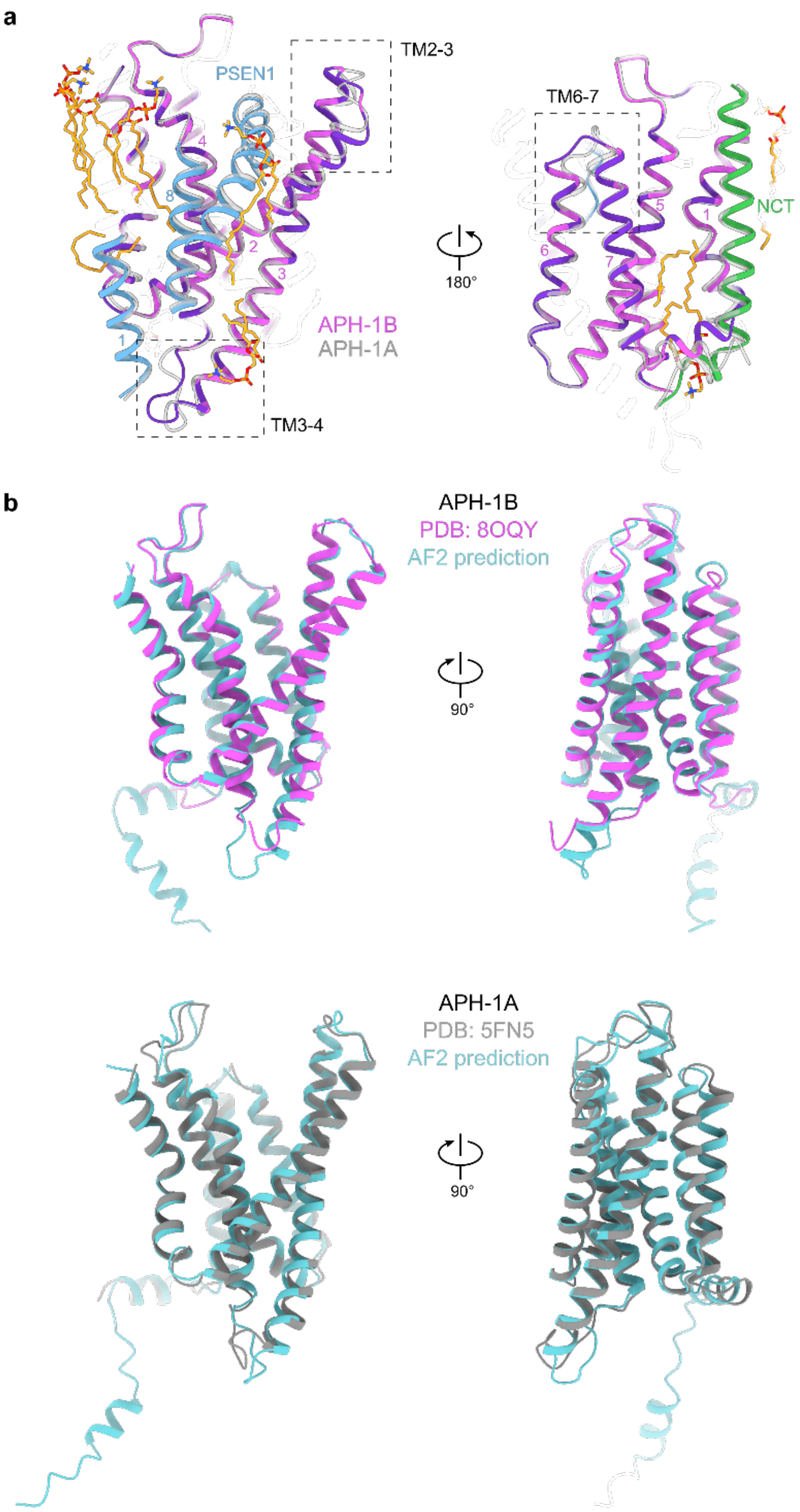


**Supplementary Figure 1. Comparison of APH-1 isoforms in substrate-bound structures with APH-1 structures predicted by AlphaFold2. a,** Structural differences between APH-1 isoforms from the GSEC1B-Aβ46 and GSEC1A-APP$_{C83}$ structures. Non-conserved APH-1B TM domain residues at the interface with PSEN1 are shown in purple. **b,** APH-1A and APH-1B structures as predicted by AF2, retrieved from UniProt (entries Q96BI3-F1-model_v4 and Q8WW43-F1-model_v4).

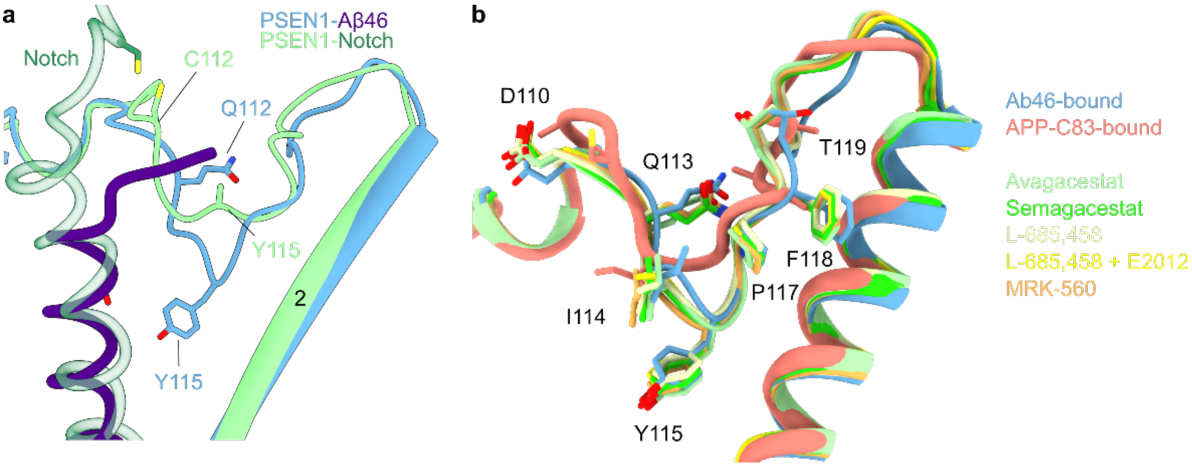


**Supplementary Figure 6. Comparison of loop1$^{PSEN1}$ conformations between GSEC structures. a,** GSEC1A-Notch (PDB:6IDF) structure is shown aligned to GSEC1B-Aβ46 structure. Side chains mutated to cysteines to form disulphide crosslinking (P9C on Notch and Q112C on PSEN1) are shown as sticks. **b,** Alignment of loop 1 from several GSEC structures solved in the presence of inhibitors[73–75]. The inhibitors used are indicated on the right and colour coded to match the structures. Loop 1 backbone is shown as cartoon, and selected residues are shown as sticks.

**Supplementary Table 1. Sequence modelled for individual GSEC subunits.** The model completeness is shown for GSEC1A, columns 1 and 3, and GSEC1B solved in this work, columns 2 and 4.

| | **GSEC1A apo (PDB: 5FN5)** | **GSEC1B apo (PDB: 8OQY)** | **GSEC1B-C83 (PDB:6IYC)** | **GSEC1B-Ab46 (PDB: 8OQZ)** |
|---|---|---|---|---|
| NCT | 34-698 (94%) | 34-699 (94%) | 34-700 (94%) | 34-610, 613-700 (94%) |
| PSEN1 | 68-108, 167-264, 379-429, 435-467 (48%) | 73-107, 160-256, 280-289, 383-467 (49%) | 73-291, 376-467 (67%) | 73-291, 376-467 (67%) |
| APH-1A | 2-244 (92%) | - | 2-244 (92%) | - |
| APH-1B | - | 2-104, 111-238 (90%) | - | 2-241 (93%) |
| PEN-2 | 2-21, 25-44, 46-101 (95%) | 2-101 (99%) | 2-101 (99%) | 2-101 (99%) |

**Supplementary Table 2. RMSD between aligned structures of GSEC1A and GSEC1B isoforms.**

| **GSEC1B PDB: 8OQY** | **GSEC1A PDB: 5FN5** | **RMSD (Å)** | **Nr of atoms** |
|---|---|---|---|
| Overall | Overall | 1.2 | 7918 |
| Nicastrin | Nicastrin | 0.9 | 4416 |
| PSEN1 | PSEN1 | 1.3 | 1353 |
| APH-1B | APH-1A | 0.9 | 1283 |
| PEN-2 | PEN-2 | 1.1 | 646 |

| **GSEC1B PDB: 8OQY** | **GSEC1B-Aβ46 PDB: 8OYZ** | **RMSD (Å)** | |
|---|---|---|---|
| Overall | Overall | 0.8 | 7365 |
| Nicastrin | Nicastrin | 0.4 | 4127 |
| PSEN1 | PSEN1 | 1.5 | 1316 |
| APH-1B | APH-1B | 0.4 | 1332 |
| PEN-2 | PEN-2 | 0.6 | 580 |

| **GSEC1B-Aβ46 PDB: 8OQZ** | **GSEC1A-C83 PDB: 6IYC** | **RMSD (Å)** | |
|---|---|---|---|
| Overall | Overall | 0.8 | 8449 |
| Nicastrin | Nicastrin | 0.5 | 4264 |
| PSEN1 | PSEN1 | 0.6 | 1823 |
| APH-1B | APH-1A | 0.7 | 1340 |
| PEN-2 | PEN-2 | 0.6 | 654 |
| **Aβ46** | APP-C83 | 1.0 | 85 |

| **GSEC1A PDB: 5FN5** | **GSEC1A-C83 PDB: 6IYC** | **RMSD (Å)** | |
|---|---|---|---|
| Overall | Overall | 1.3 | 7808 |
| Nicastrin | Nicastrin | 0.8 | 4457 |
| PSEN1 | PSEN1 | 2.1 | 1431 |
| APH-1A | APH-1A | 0.9 | 1530 |
| PEN-2 | PEN-2 | 1.2 | 694 |

**Supplementary Table 3. List of APH-1 residues located at the interface with PSEN1 subunit.**

| APH-1A | APH-1B | Change APH-1A -> APH-1B | PSEN1 sidechains within 4.5 Å | |
|---|---|---|---|---|
| | | | APH-1A | APH-1B |
| V32 | I32 | Longer | F86 | F86 |
| V36 | I36 | Longer | F86, L415, I416, C419, L420 | F86, V412, L415, I416 |
| V51 | L51 | Longer | L452 | L452 |
| R62 | N62 | Positive -> polar | Q459 | - |
| Y69 | K69 | Aromatic -> positive | - | F465 |
| L104 | I104 | - | E72 | A79 |
| D107 | G107 | Positive -> no sidechain | L73, K76 | - |
| I128 | M127 | Longer | V412 | V412 |
| I135 | V134 | Shorter | W404, N405 | W404 |
| I137 | T136 | Hydrophobic -> polar | A461 | A461 |
| Y156 | F155 | Polar -> hydrophobic | Q464 | Q464 |
| T159 | Y158 | Liner -> aromatic | Q464 | Q464, Y466 |
| L163 | M162 | Longer | Y466 | Y466 |
| T200 | V199 | Polar -> hydrophobic | - | - |
| N207 | S206 | Shorter | Q464, F465, Y466 | - |

**Supplementary Video 1. Conformational changes between GSEC1B in nanodiscs and GSEC1A in amphipols.**

**Supplementary Video 2. Conformational changes between GSEC1B in the apo and Aβ46-bound state.**

**Supplementary Video 3. Conformational changes between GSEC1B-Aβ46 and GSEC1A-C83.**
